# Genome-wide analysis clarifies the population genetic structure of wild Gilthead Sea Bream (*Sparus aurata*)

**DOI:** 10.1101/2020.07.06.189241

**Authors:** Francesco Maroso, Konstantinos Gkagkavouzis, Sabina De Innocentiis, Jasmien Hillen, Fernanda do Prado, Nikoleta Karaiskou, John Bernard Taggart, Adrian Carr, Einar Nielsen, the Aquatrace consortium, Alexandros Triantafyllidis, Luca Bargelloni

**Affiliations:** Department of Comparative Biomedicine and Food Science, University of Padua, 35020 Legnaro (PD), Italy; Department of Genetics, Development and Molecular Biology, School of Biology, Aristotle University of Thessaloniki, 54 124 Thessaloniki, Macedonia, Greece; ISPRA, Institute for Environmental Protection and Research, 00144 Roma, Italy; Laboratory of Biodiversity and Evolutionary Genomics, University of Leuven, 3000 Leuven, Belgium; Department of Biological Sciences, São Paulo State University, 01049-010 Bauru, Brazil; Institute of Aquaculture, University of Stirling, FK9 4LA Stirling, United Kingdom; Fios Genomics Ltd, EH16 4UX Edinburgh, United Kingdom; Section for Population Ecology and Genetics, National Institute of Aquatic Resources, Technical University of Denmark, DK-8600 Silkeborg, Denmark

## Abstract

Gilthead sea bream is an important target for both recreational and commercial fishing in Europe, where it is also one of the most important cultured fish. Its distribution range goes from the Mediterranean to the African and European coasts of the North-East Atlantic. So far, the genetic structure of this species in the wild has been studied with microsatellite DNA, but the pattern of differentiation could not be fully clarified. In this study, almost 1000 wild sea bream from 23 locations in the Mediterranean Sea and Atlantic ocean where genotyped at 1159 SNP markers, of which 18 potentially under selection. Neutral markers suggested the presence of a weak subdivision into three genetic clusters: Atlantic, West and East Mediterranean. This last group could be further subdivided into an Ionian/Adriatic and an Aegean group using outlier markers. Seascape analysis suggested that this differentiation was mainly due to difference in salinity, and this was also supported by preliminary genomic functional analysis. These results are of fundamental importance for the development of proper management of this species in the wild and are a first step toward the study of the potential genetic impact of the sea bream aquaculture industry.

## 1. Introduction

Compared to studies of terrestrial animals, exploring population genetic structure in marine species can be challenging. This is particularly true for fish, which are among the most motile marine organisms, often throughout their entire life cycle. New genomic approaches like Next Generation Sequencing (NGS) and Restriction site Associated DNA (RAD) have already proven to be effective in detection of hidden population structure and to better define unclear genetic subdivision and differentiation in fish [1–6]. Higher number of markers also increases the chances of detecting genomic regions under selective pressure [2,7,8] allowing the investigation of demographic *vs* environmental causes of genetic differentiation among populations. Such molecular markers can also be applied for the geographical traceability of wild samples, to be implemented in actions against illegal, unreported and unregulated (IUU) fishing as well as to meet the increased interest of customers about food origin. In a wider context, they also enable detailed comparison of wild and farmed populations to predict the potential impact of aquaculture on natural populations, a recommended practice since the early ‘90s [9–14].

Sea bream is a protandric hermaphrodite, demersal species living in warm coastal and euryhaline waters of the Mediterranean Sea and North-East Atlantic Ocean [15]. It is highly appreciated for the quality of its flesh and it is an important target for both commercial and recreational fishing. Capture fisheries have provided almost constant production since the ’60s (around 8000 tons per year) and aquaculture production has increased from the early ’90s and reached 186,000 tons in 2016 [16]. Following the expansion of marine cage culture, concerns have arisen regarding the potential effects of farm escapes on the natural populations [14,17].

Current knowledge of the genetics of gilthead sea bream is scarce and fragmented for wild populations. Previous studies of the natural genetic structuring along sea bream distribution range did not provide a consistent scenario, and while some surveys report an absence of genetic differentiation among basins [18], others reported subtle genetic structure or population subdivision even at small geographical scale [19–22]; also, it cannot be excluded that aquaculture practices, also those involving other species (e.g. tunas or shellfish) have an effect in shaping the population genetic structure of the species [23]. Anyhow, these studies were mainly based on markers (e.g. microsatellites, mitochondrial DNA) which are nowadays outperformed by SNPs. The lack of a consensus on the genetic distinction between populations has also hindered the development of genetic traceability tools, which are based on the definition of reference units.

In the current study we used RAD to identify and score a large, genome wide panel of more than 1000 SNPs in a geographically diverse set of sea bream samples (more than 1000 specimens from 23 locations), in order to better characterize the genetic variability and the differentiation across North-East Atlantic and Mediterranean basins. The effect of environmental variables, as well as demographic processes, are assessed and discussed in context of life history of the species and previous knowledge of sea bream’s population genetics. The data provide a valuable baseline for future management of the wild stocks and for the assessment of potential impact of the aquaculture on the genetic of the species.

## 2. Material and Methods

A total of 956 wild individuals from 23 different locations were sampled (Figure 1 and Table 1). For three populations (GRE-6, GRE-7 and GRE-9) temporal replicates were also available. These older samples were compared with more recent ones from the same area but then excluded from further analysis (i.e. cluster analysis and outlier detection). Samples consisted of either fin clips or muscle tissue, preserved in 95% ethanol as soon as possible after sampling. Genomic DNA was extracted using the commercial kit Invisorb^®^ Spin Tissue Mini Kit (Invitek, STRATEC Biomedical, 242 Germany) or the SSTNE buffer, a modified TNE buffer including spermidine and spermine [24], that allowed a more efficient, though more time consuming, extraction of samples that failed with the commercial kit.

**Fig. 1.**
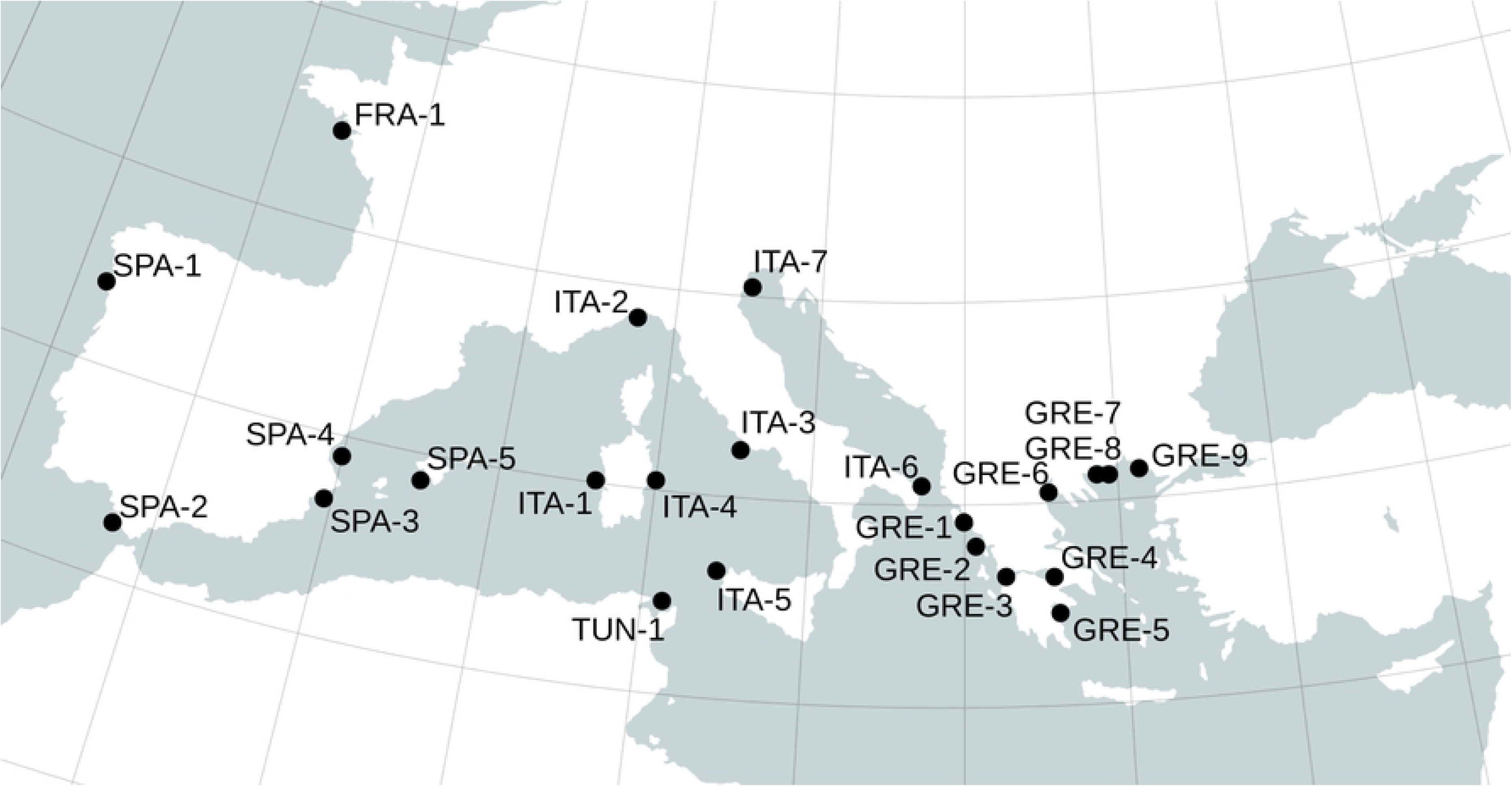
Map of sampling locations.

**Table 1.**
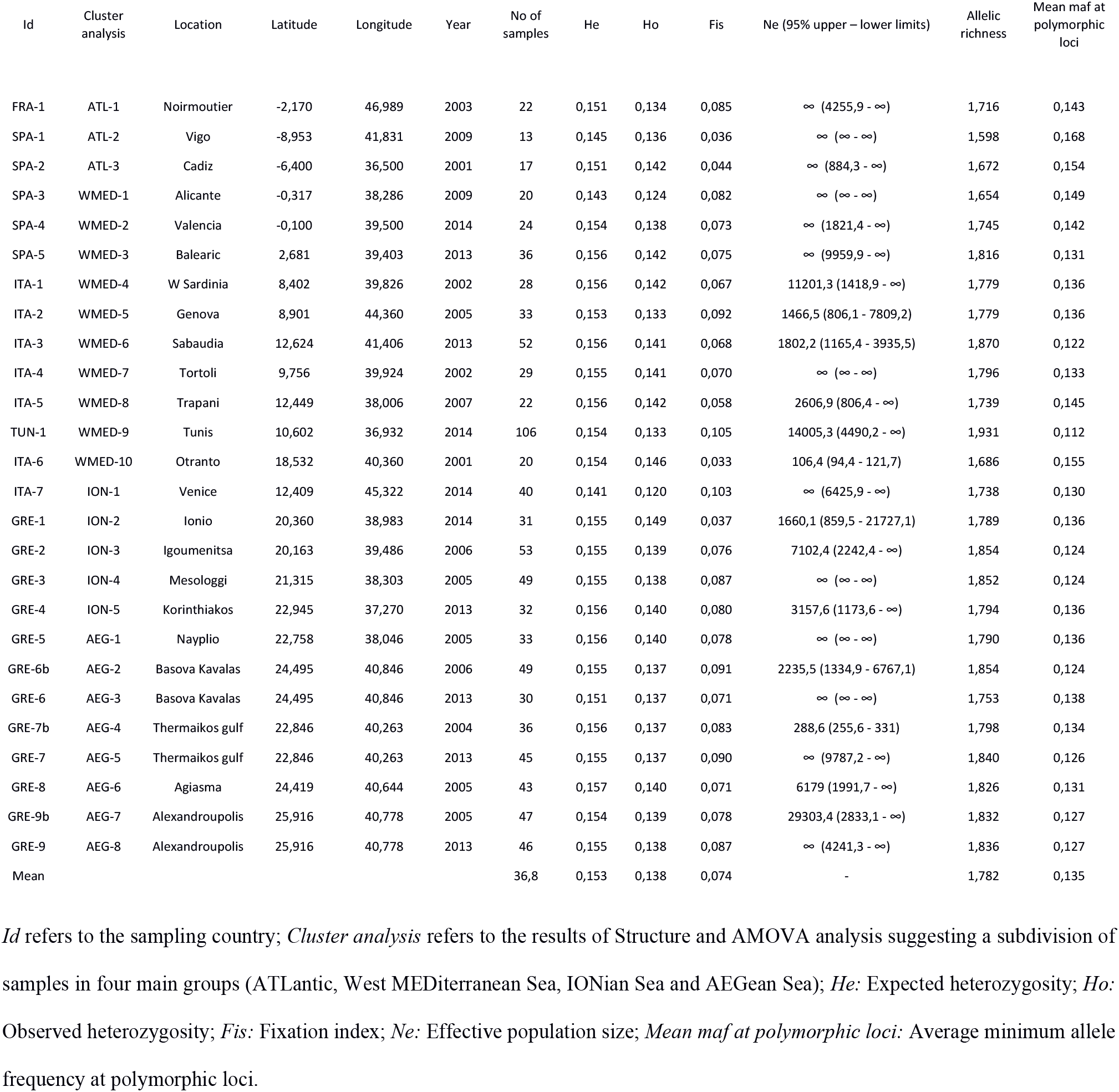
Sampling groups and basic statistics.

| Id | Cluster analysis | Location | Latitude | Longitude | Year | No of samples | He | Ho | Fis | Ne (95% upper – lower limits) | Allelic richness | Mean maf at polymorphic loci |
| --- | --- | --- | --- | --- | --- | --- | --- | --- | --- | --- | --- | --- |
| FRA-1 | ATL-1 | Noirmoutier | -2,170 | 46,989 | 2003 | 22 | 0,151 | 0,134 | 0,085 | ∞ (4255,9 - ∞) | 1,716 | 0,143 |
| SPA-1 | ATL-2 | Vigo | -8,953 | 41,831 | 2009 | 13 | 0,145 | 0,136 | 0,036 | ∞ (∞ - ∞) | 1,598 | 0,168 |
| SPA-2 | ATL-3 | Cadiz | -6,400 | 36,500 | 2001 | 17 | 0,151 | 0,142 | 0,044 | ∞ (884,3 - ∞) | 1,672 | 0,154 |
| SPA-3 | WMED-1 | Alicante | -0,317 | 38,286 | 2009 | 20 | 0,143 | 0,124 | 0,082 | ∞ (∞ - ∞) | 1,654 | 0,149 |
| SPA-4 | WMED-2 | Valencia | -0,100 | 39,500 | 2014 | 24 | 0,154 | 0,138 | 0,073 | ∞ (1821,4 - ∞) | 1,745 | 0,142 |
| SPA-5 | WMED-3 | Balearic | 2,681 | 39,403 | 2013 | 36 | 0,156 | 0,142 | 0,075 | ∞ (9959,9 - ∞) | 1,816 | 0,131 |
| ITA-1 | WMED-4 | W Sardinia | 8,402 | 39,826 | 2002 | 28 | 0,156 | 0,142 | 0,067 | 11201,3 (1418,9 - ∞) | 1,779 | 0,136 |
| ITA-2 | WMED-5 | Genova | 8,901 | 44,360 | 2005 | 33 | 0,153 | 0,133 | 0,092 | 1466,5 (806,1 - 7809,2) | 1,779 | 0,136 |
| ITA-3 | WMED-6 | Sabaudia | 12,624 | 41,406 | 2013 | 52 | 0,156 | 0,141 | 0,068 | 1802,2 (1165,4 - 3935,5) | 1,870 | 0,122 |
| ITA-4 | WMED-7 | Tortoli | 9,756 | 39,924 | 2002 | 29 | 0,155 | 0,141 | 0,070 | ∞ (∞ - ∞) | 1,796 | 0,133 |
| ITA-5 | WMED-8 | Trapani | 12,449 | 38,006 | 2007 | 22 | 0,156 | 0,142 | 0,058 | 2606,9 (806,4 - ∞) | 1,739 | 0,145 |
| TUN-1 | WMED-9 | Tunis | 10,602 | 36,932 | 2014 | 106 | 0,154 | 0,133 | 0,105 | 14005,3 (4490,2 - ∞) | 1,931 | 0,112 |
| ITA-6 | WMED-10 | Otranto | 18,532 | 40,360 | 2001 | 20 | 0,154 | 0,146 | 0,033 | 106,4 (94,4 - 121,7) | 1,686 | 0,155 |
| ITA-7 | ION-1 | Venice | 12,409 | 45,322 | 2014 | 40 | 0,141 | 0,120 | 0,103 | ∞ (6425,9 - ∞) | 1,738 | 0,130 |
| GRE-1 | ION-2 | Ionio | 20,360 | 38,983 | 2014 | 31 | 0,155 | 0,149 | 0,037 | 1660,1 (859,5 - 21727,1) | 1,789 | 0,136 |
| GRE-2 | ION-3 | Igoumenitsa | 20,163 | 39,486 | 2006 | 53 | 0,155 | 0,139 | 0,076 | 7102,4 (2242,4 - ∞) | 1,854 | 0,124 |
| GRE-3 | ION-4 | Mesologgi | 21,315 | 38,303 | 2005 | 49 | 0,155 | 0,138 | 0,087 | ∞ (∞ - ∞) | 1,852 | 0,124 |
| GRE-4 | ION-5 | Korinthiakos | 22,945 | 37,270 | 2013 | 32 | 0,156 | 0,140 | 0,080 | 3157,6 (1173,6 - ∞) | 1,794 | 0,136 |
| GRE-5 | AEG-1 | Nayplio | 22,758 | 38,046 | 2005 | 33 | 0,156 | 0,140 | 0,078 | ∞ (∞ - ∞) | 1,790 | 0,136 |
| GRE-6b | AEG-2 | Basova Kavalas | 24,495 | 40,846 | 2006 | 49 | 0,155 | 0,137 | 0,091 | 2235,5 (1334,9 - 6767,1) | 1,854 | 0,124 |
| GRE-6 | AEG-3 | Basova Kavalas | 24,495 | 40,846 | 2013 | 30 | 0,151 | 0,137 | 0,071 | ∞ (∞ - ∞) | 1,753 | 0,138 |
| GRE-7b | AEG-4 | Thermaikos gulf | 22,846 | 40,263 | 2004 | 36 | 0,156 | 0,137 | 0,083 | 288,6 (255,6 - 331) | 1,798 | 0,134 |
| GRE-7 | AEG-5 | Thermaikos gulf | 22,846 | 40,263 | 2013 | 45 | 0,155 | 0,137 | 0,090 | ∞ (9787,2 - ∞) | 1,840 | 0,126 |
| GRE-8 | AEG-6 | Agiasma | 24,419 | 40,644 | 2005 | 43 | 0,157 | 0,140 | 0,071 | 6179 (1991,7 - ∞) | 1,826 | 0,131 |
| GRE-9b | AEG-7 | Alexandroupolis | 25,916 | 40,778 | 2005 | 47 | 0,154 | 0,139 | 0,078 | 29303,4 (2833,1 - ∞) | 1,832 | 0,127 |
| GRE-9 | AEG-8 | Alexandroupolis | 25,916 | 40,778 | 2013 | 46 | 0,155 | 0,138 | 0,087 | ∞ (4241,3 - ∞) | 1,836 | 0,127 |
| Mean |  |  |  |  |  | 36,8 | 0,153 | 0,138 | 0,074 | - | 1,782 | 0,135 |

*Id* refers to the sampling country; *Cluster analysis* refers to the results of Structure and AMOVA analysis suggesting a subdivision of
samples in four main groups (ATLantic, West MEDiterranean Sea, IONian Sea and AEGean Sea); *He*: Expected heterozygosity; *Ho*:
Observed heterozygosity; *Fis*: Fixation index; *Ne*: Effective population size; *Mean maf at polymorphic loci*: Average minimum allele
frequency at polymorphic loci.

Multiple ddRAD libraries were prepared, each including 144 samples, splitting samples from the same population in different libraries in order to avoid confounding library-specific biases. Library preparation protocol followed the original one of Peterson et al. [25], with some modifications that facilitate the screening of a larger number of individuals [26,27] (Supplementary Material S1).

Raw reads were checked for quality using FASTQC [28]. Then, reads missing valid restriction site were discarded and barcodes were searched (allowing up to one error) for demultiplexing. Barcodes were removed and the remaining sequences were trimmed to 90 bp length. Four bases at the end of each read were cut in order to increase the number of reads passing the filter and to obtain higher coverage at the end of the genotyping process. Reads with one or more uncalled bases were filtered out, as well as reads with 11 or more consecutive bases with an average quality score less than 20 (1% error rate). If a sample was sequenced on more than one lane, reads were combined into a single file before processing. Stacks 1.3 [29,30] was used to cluster reads into consensus tags and call high-quality SNPs. Stacks' *de-novo* pipeline was run with minimum depth of coverage to call a stack (-m) set to four; maximum number of differences between stacks to be considered as the same tag in *ustacks* and *cstacks* (-M and -n, respectively) set to seven; SNP calling model was set to ‘bounded’. Correction module *rxstacks* was run after the analysis to correct genotypes based on population-wide information (refer to Stacks’ website for details about the pipeline). Since including all samples in the catalogue construction would be prohibitively time-consuming with the version of Stacks used, 500 samples were selected for this step, including those with a higher number of reads from each population, in order to have all of them represented. Finally, SNPs were filtered out if scored in less than 80% of the analyzed samples and when Minor Allele Frequency (MAF) was lower than 0.5%. Similarly, samples were filtered in order to retain only those genotyped at more than 80% of the markers.

Four samples were replicated 12 to 13 times in different libraries in order to assess genotyping precision. For each locus, genotypes of replicates were compared using the most frequent one as a reference and accounting the number of mismatches across all replicates.

GenAlEx 6.501 [31] was used to calculate expected (He) and observed (Ho) heterozygosity and allelic richness of polymorphic markers (AR). Deviation from Hardy-Weinberg equilibrium (excess or defect of heterozygotes) was tested for each locus and for each population using Genepop 4.6 [32]. We tested for unusually high LD between pairs of loci using r^2^estimator implemented in Plink 1.9 [33], parsing all pairs of loci. F_ST_ matrices were calculated with Arlequin 3.5.2.1 [34] using 50,000 permutations to test for significance. AMOVA was also performed using Arlequin, to test the clustering suggested by other analysis.

Two different approaches were used to summarize and visualize the genetic relationship between groups: the model-based clustering method implemented in Structure [35] and Discriminant Analysis of Principal Components (DAPC) as implemented in the *adegenet* R package [36,37]. Structure 2.3.4 was run through Parallel Structure [38] to allow faster and more efficient parallel running using different *k* values and replicates of each *k* value. We ran the algorithm using the sampling location as *a-priori* information. The analysis was run with *k* ranging from one to ten, each repeated five times to allow evaluation of the likelihood of different simulated number of ancestral clusters. Burn in (BI) was set to 50,000 and the number of iterations (IT) to 100,000. Results from different runs were collated and most likely k values were detected using the Evanno’s method implemented in STRUCTURE HARVESTER [39]. A further Structure run was carried out with the most likely *k* value, using 100,000 burn-in cycles and 300,000 iterations. DAPC was carried out using R’s package *adegenet* [37]. To avoid the effect of retaining too many principal components (PC), which would discriminate better the sampled individuals, whilst performing poorly with newly sampled ones, repeated cross-validation was used to select the best number of SNPs and to obtain a trade-off between stability and power of discrimination.

To evaluate the level of genetic diversity of the populations analyzed we estimated Effective population Size (Ne) of the natural populations using a single sample method based on the increased level of Linkage Disequilibrium between loci, that arise when populations with low Ne are sampled. The algorithm is implemented in NeEstimator 2.01 [40,41]. Ne were estimated using only putatively neutral SNPs (see below) with minor allele frequency (MAF) >1%. Pairwise genetic relatedness between individuals was calculated with Coancestry 1.0 [42].

One of the most interesting and useful advantages of genome-wide genotyping is the increased chance of finding genetic regions (i.e. loci) potentially under natural selection. These markers can be used to link genetic and phenotypic traits selected in a particular environment. In this work, we used the Bayesian approach implemented in Bayescan 2.1 [43,44] and Arlequin [34]. A combination of different methods is indeed advised to obtain reliable information from the data [45]. A similar analysis was carried out within Atlantic and Mediterranean basins and with samples divided into four groups, as suggested by clustering analysis (see Results). Both programs were run with default parameters and finally, an outlier panel (OL) was defined selecting only loci detected by both methods using stringent threshold (Log10 (PO) > 1.5 for Bayescan and p < 0.05 for Arlequin) to avoid including false positives. The neutral dataset was defined excluding the potential OL suggested by at least one of the methods applied. To understand how environmental factors and spatial variables shaped genetic diversity across sea bream distribution range, we applied redundancy analysis (RDA) using R's VEGAN library [46–48]. Environmental data was extracted from SeaDataNet portal (http://www.seadatanet.org/) and included winter and summer temperature and salinity values at the surface and at 20 m, as these represent functional proxies of more complex environmental variation (Supplementary Material S2 Table “ENVIRON”). We referred to the geographic coordinates as close as possible to actual sampling locations for which data were available. The analysis was carried out using the entire SNP dataset and the reduced neutral and outlier datasets. Since the neutral and the full datasets provided similar results, only the former will be presented. The significance of the relationship between environmental and genetic data was tested with permutation tests and variance explained by different independent factor was taken into account. The panels of explanatory variables were reduced by automatic forward selection based on significant variable criteria and the total proportion of genetic variation was recalculated accordingly. Variation in environmental data values was also compared to variation in allele frequency to look for a correlation between environmental variable values and allele frequency. For this purpose, we used BayEnv 2 [49,50]. Four separated analyses were run to calculate correlation values and locus/variable specific Bayes Factors (BFs) were averaged to obtain more reliable results. RADtags containing potential outlier loci, as well as loci whose allele frequency was significantly correlated with environmental factors were located in the sea bream genome [51] using BLAST to identify the presence of genes in the regions adjacent to the markers detected. Genes annotations were extracted from the regions flanking each OL, spanning 50k bp up and downstream from tag location, using a custom-made Perl script. To detect variation in features under multigenic control, enrichment analysis was carried out for the list of genes extracted with Blast2GO, using the sea bream genome annotation as a reference to identify over or under-represented biological processes, cellular components or molecular functions. Results were reported only for features with p-value < 0.05.

## 3. Results

### 3.1 Almost one thousand wild sea breams consistently genotyped at 1159 high quality SNP

A total of 767.1 M read pairs were successfully sequenced, with an average of 804,463 ± 505,000 reads per individual (range 97 – 3,104,802 reads). After filtering, an average of 72.2% ± 6% of reads was retained. The initial number of called SNPs was 11,662, included in a total of 216,713 tags. After filtering out low-quality markers, 1165 SNPs (10.6%) were retained. After filtering, the level of data missing per sample ranged from 0% to 19.9%, with an average of 5.4%. Of 30,290 tests for departure from H-W equilibrium carried out, after sequential Bonferroni correction, two loci showed significant deviation from equilibrium (both for an excess of heterozygosity) in more than half of the natural populations and were excluded. A total of 675,703 tests for LD were carried out and four loci pairs showed r^2^values higher than 0.7 and for each pair, the locus with lower missing data was retained. Remaining 1159 SNPs were used for subsequent analysis (Supplementary Material S3). The mismatch rate between replicates at 1159 filtered loci ranged from 3.4% to 5.8 %, with an average of 4.0%. Mean number of alleles per locus (Na) was 1.782, ranging from 1.598 (SPA-1) to 1.931 (TUN-1). Mean expected and observed heterozygosity were 0.153 (range 0.141 – 0.157) and 0.138 (range 0.120 – 0.149), respectively (Table1). All these parameters tend to be lower in the Atlantic populations. Minor Allele Frequency (MAF) ranged from 0.112 to 0.168, with an average of 0.135. We found a trend of decreased Na, He and Ho and increased MAF in Atlantic samples, that was also correlated to an average lower number of samples for this area. Despite this, Effective Population Size (Ne) for all Atlantic samples were very high ('Infinite'), whereas Mediterranean samples showed values ranging from 106.4 (ITA-6) to 'Infinite' (ten samples). Nevertheless, ranges of confidence varied substantially especially for smaller samples, which suggests caution when interpreting this data. Inbreeding index varied from 0.033 to 0.105 and averaged 0.074. No general trend was found for this parameter when comparing Atlantic and Mediterranean samples. Relatedness analysis gave clues for the presence of potential full-sib pairs within SPA-2 sample (relatedness > 0.375). As a consequence, four individuals that were part of highly related pairs were eliminated from the dataset to avoid biases in the clustering analysis. Genetic differentiation between time replicates was not significant. Heterozygosity was also similar in time replicates, but for Basova Kavalas and Thermaikos Gulf recent samples showed significantly higher Ne.

### 3.2 Four genetic clusters identified in sea bream distribution area

Comprehensively, the results obtained with the full SNPs dataset (representing the combination of demographic and selective factors) suggested a subdivision of sea bream in four genetic clusters: Atlantic (ATL), West Mediterranean (WMED), Ionian/Adriatic seas (ION) and the Aegean Sea (AEG), the strongest differentiation being between Atlantic and Mediterranean samples. The Evanno’s method suggested a most likely number of four clusters and all runs at this number of ancestral populations showed the same cluster pattern. Samples showed a high level of admixture, but samples from different basins differed in the proportions of each component. Within-basin differences were much lower (Figure 2). Samples from Northern and Southern Adriatic Sea (i.e. ION01/ITA7 and WMED10/ITA6, respectively) showed no genetic differentiation (F_ST_= 0.0000), however under Structure’s Bayesian analysis they clustered separately: with ION and WMED populations, respectively. AMOVA analysis further confirmed the goodness of this subdivision, as the proportion of genetic variability among basins was statistically significant, while the variability within groups was not statistically significant (Table 2).

**Fig. 2.**
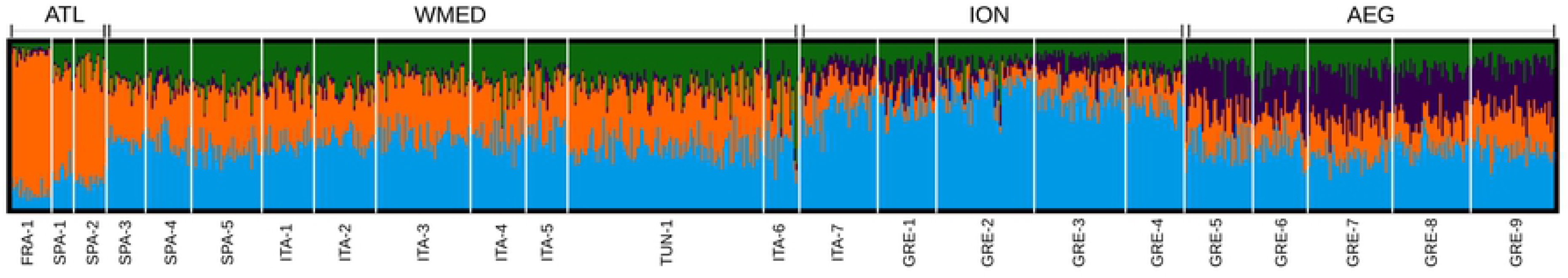
Result of Structure clustering analysis for the most likely value for the number of cluster (*k* = 4). Labels under the graph indicate the sampling groups. Labels above the graph indicate the genetic clusters suggested by the analysis and supported by AMOVA.

**Table 2.** Results of the AMOVA analysis with sample subdivided according to the clustering analysis.

| Source of Variation | Degrees of freedom | Sum of squares | Variance components | Fixation indices | Percentage of Variation | P-value |
| --- | --- | --- | --- | --- | --- | --- |
| Among groups | 3 | 433.523 | 0.29020 | 0.00363 | 0.36 | <0.01 |
| Among populations within groups | 19 | 732.743 | -0.66080 | -0.00830 | -0.83 | ns |
| Among individuals within populations | 801 | 68161.210 | 4.82579 | 0.06012 | 6.04 | <0.01 |
| Within individuals | 824 | 62165.500 | 75.44357 | 0.05576 | 94.42 | <0.01 |
| Total | 1647 | 131492.976 | 79.89876 |  | - | - |

DAPC was based on 150 PCs, after cross-validation analysis. Differentiation signal was weak, with the first axis roughly matching the geographical West to the East distribution of the samples. Along the second axis of the DAPC ATL and AEG clusters showed some level of genetic similarity when compared to the two remaining clusters of Mediterranean (Supplementary Material S4).

F_ST_ values were in general low (average F_ST_=0.0031) with a trend of increased values in the comparisons between Atlantic and Mediterranean (Supplementary Material S5). Within the Mediterranean, values tend to increase and be more significant in the comparisons between samples from the Western and Eastern Mediterranean (Ionian and Aegean basins). When samples are subdivided in the proposed four groups, F_ST_ values were low but highly significant (p<0.001) and ranging from 0.0015 in the comparison between WMED and AEG to 0.0070 in the comparison between the ION and ATL clusters. No loci showed private alleles across wild populations, but one locus (7704_13) showed a private allele in WMED group, genotyped in nine heterozygous individuals, when compared to the other groups.

### 3.3 Dissecting genetic differentiation: Atlantic-Mediterranean differentiation at neutral loci and signs of convergent selection at outlier loci

Bayescan detected 20 loci with log10(PO) > 1.5; Arlequin detected 28 potential diverging outliers at p < 0.05. A total of 16 loci (1.3% of the entire dataset) shared by the two methods were selected to create the ‘outlier dataset’ (OL, Supplementary Material S3). Two more loci were added as they were detected the analysis with samples divided into four clusters (see above). OL dataset was subsequently used for analysis focused at understanding the “functional” divergence between populations. Putative neutral loci dataset consisted of 1126 remaining loci, after excluding potential outliers identified by at least one of the approaches used.

An additional SNP (8727_39) was detected by both approaches but was not included in the subsequent analysis due to unclear results obtained when it was genotyped with single SNP allele-specific assay. On the other side, with the same technique all the other OL SNPs were confirmed (data not shown).

BayEnv detected 24 loci, of which six were among those identified by the joint analysis carried out with Bayescan and Arlequin (Supplementary Material S3).

OL allele frequencies showed different behavior when moving from West to East populations: some followed a gradual variation (e.g. 2689_65); some other showed more abrupt changes at basin boundaries (e.g. 12615_64); finally, as expected, OL identified by BayEnv showed patterns of allele frequency that overlapped with environmental variables (e.g. 13310_71). This is true also for markers not previously identified by other OL methods (e.g. 13776_28) (Supplementary Material S6).

The genetic structure resulting from the panel of putative neutral loci was weaker than using the entire dataset, especially within the Mediterranean. F_ST_ values were rarely significant and mostly in the comparisons between Atlantic samples and samples from Ionian and Aegean seas (Supplementary Material S7). Structure clustering analysis at the most likely value of k=2 (according to the Evanno's method) showed high admixture with some level of differentiation in cluster representation between ATL and WMED samples. ION and AEG samples looked similar to each other and slightly differentiated from the remaining two groups (Supplementary Material S8).

With the OL panel, not only the F_ST_ differentiation generally increased, but also the presence of within basin differentiation emerged (Supplementary Material S7). Northern Atlantic sample ATL1/FRA-1 was differentiated from more southern samples ATL2/SPA-1 and ATL3/SPA-2. In general, ATL1/FRA-1 is the most differentiated sample with F_ST_ ranging up to 0.1751, in the comparison with samples ION3/GRE-2. WMED is the only group that is highly homogeneous at outlier loci. The most striking difference in the result obtained with this dataset compared to the neutral one is the strong differentiation between Ionian and Aegean samples, reflected in F_ST_ values and Structure results at the most probable k=3 (Supplementary material S9).

Genetic differentiation at outlier loci reflected environmental differentiation between localities, as confirmed by SEASCAPE analysis, that indicated Longitude and winter surface salinity (S_0_W) as significant explanatory variables. Climate explained a total of 54% of the variance in the data, while geography explained 14%. The joint effect of the two sets of variables explained 32% of the total variance. The plot indicated that Longitude had the strongest effect on the first axis, while winter surface salinity was correlated with the second axis of the PCA (Figure 3). Along this axis, ATL samples are roughly grouped in the lower half of the graph and closer to AEG and ION-1/ITA-7 samples, reflecting differences in salinity values. On the other hand, ION and ATL samples showed the strongest pairwise differences both in terms of environmental variables and also in terms of differentiation at OL loci. OL were scattered in 16 of the 24 chromosomes of the species. Eighteen of them were located inside genes, 11 of which were among the markers indicated by BayEnv as correlated with environmental variables. Fourteen of these tags fell inside a Coding DNA Sequence (CDS) regions (Supplementary Material S3). The enrichment analysis suggested a significant over-representation of genes involved in the biological processes “kinetochore assembly” and “kinetochore organization” (GO:0051382 and GO:0051383) and “endoplasmic reticulum calcium ion homeostasis” (GO:0032469). In addition, pathways related to the expression of cellular membrane components seem to be involved and many “transferase” molecular functions (Supplementary Material S10).

**Figure 3.**
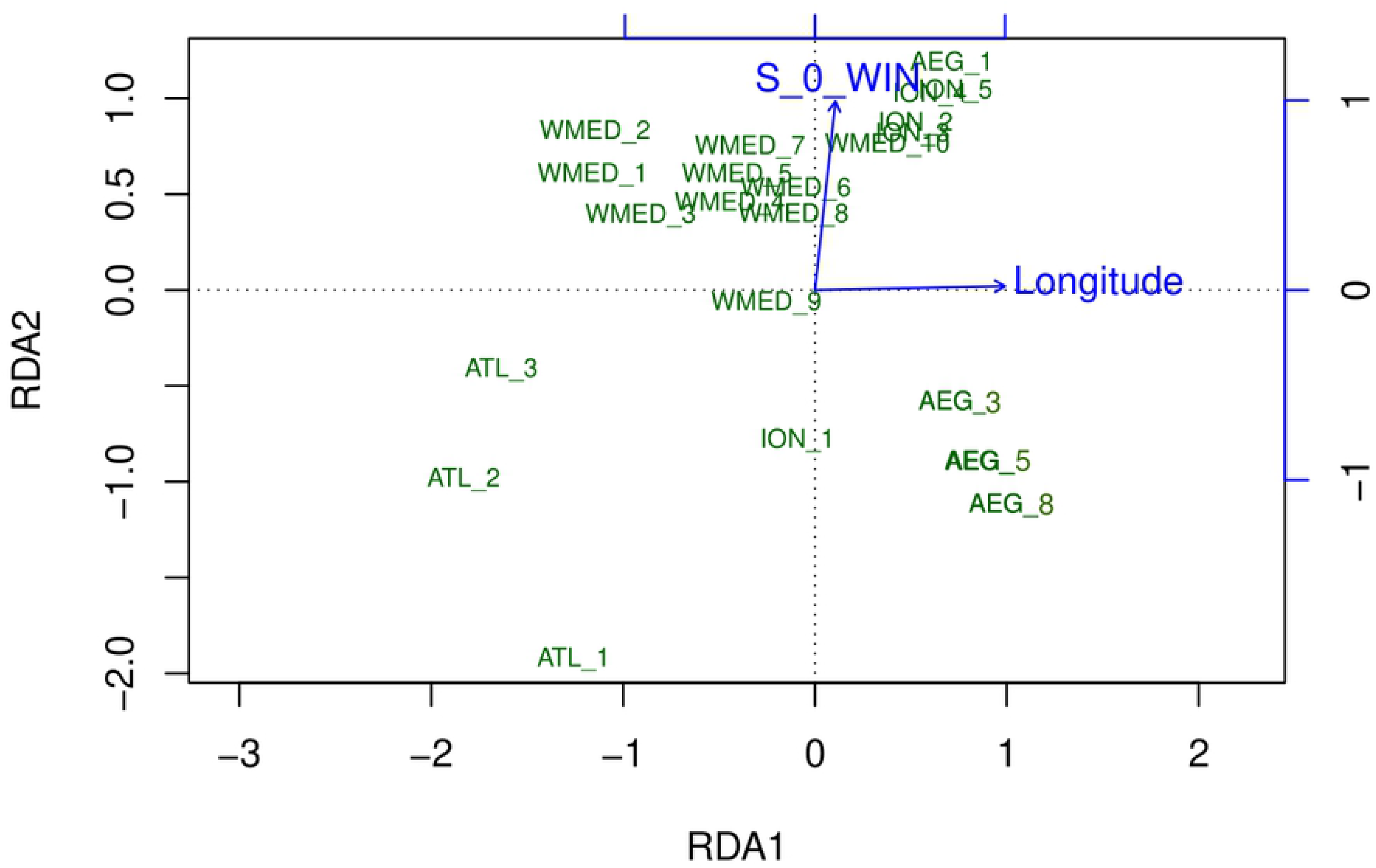
Scatterplot of the redundancy analysis performed using outlier SNP dataset. Sampling groups are labeled according to the genetic clusters identified with Structure and AMOVA and plotted according to the most explicative axes. Longitude and Surface winter salinity (S_0_WIN) were the environmental variables that resulted significantly related with genetic data.

## 4. Discussion

The genetic differentiation found is likely due to the joint effect of two main factors: (i) a demographic factor derived by long term reduced connectivity between populations; (ii) selection for the environment (i.e. short-term effect).

### 4.1 The neutral genetic structure

F_ST_ levels, according to the full SNP dataset, are lower than those usually found in fish (F_ST_=0.062 [52]) and agree with previous studies of the sea bream [18,19]. Previously, a broad geographical range analysis of gilthead seabream was performed only with allozymes and microsatellites (i.e. putatively neutral markers) by Alarcón et al. [18]. The authors concluded that the structuring pattern could not be associated with geographic nor oceanographic known factors. In our work, more sensitive approaches were used to uncover subtle genetic structure. Despite pairwise genetic differentiation was significant only in a few cases, other approaches (i.e. Structure, DAPC and AMOVA) pointed to a significant subdivision of the species in three main basins: the Atlantic, the West Mediterranean and the East Mediterranean, including the Ionian and Aegean seas.

The presence of two major oceanographic fronts separating these basins, namely the Almeria-Oran front and the Sicily Channel front, explain this pattern. In addition, the Otranto Strait front effect could also explain the differentiation between WMED10/ITA-6 and the Ionian samples. A similar result was also found by Carreras et al. [53] analyzing another coastal species: East Atlantic peacock wrasse populations. In their study, authors also found a connection between Ionian and Adriatic samples along the East coast of these basins. The same oceanographic features could explain the similarity between Ionian and North Adriatic (ION1/ITA-7) samples. These patterns could also be influenced by a stronger dispersal along the coast rather than across the opposite stretch of the sea, considering that sea bream prefers shallower water [54].

Signs of differentiation between Atlantic and Mediterranean basins were already found using other typically neutral markers (i.e. microsatellites in [19]), and significant differentiation was also found between Atlantic samples from north and south coasts of Spain. In our study, this structure was reflected by loci under selection and may be the result of an evolutionary response to the different environment. Atlantic samples collected closer to the Mediterranean (ATL3/SPA-2) looked more strongly differentiated from Mediterranean samples than ATL1/FRA-1, despite the location. A similar result was found by Alarcón et al. [18], using allozymes and microsatellites. In their study, the most differentiated population was in fact from the Atlantic south coast of Spain. Differentiation within the Mediterranean was previously found by other authors [22,55,56]. These works suggested a differentiation of the Adriatic basin and a reduced gene flow through Strait of Sicily. Low level of differentiation within basin, coupled with some signs of differentiation across fronts are expected for this species. Indeed, given the long period of larval dispersion of seabream (>30 days) [57] and its pelagic/benthic vagile lifestyle, retention rate within basins is expected to be low, with fronts and currents having a dominant effect on shaping the neutral genetic structure. Other studies of sea bream detected some level of differentiation between samples from the north and south West Mediterranean Sea [58]. In our case, the only samples from the north coast of Africa come from Tunisia and did not look differentiated from other West Mediterranean samples.

The lack of previous genetic analysis of sea bream wild populations based on SNPs does not allow comparison in term of genetic diversity, but the value of genetic diversity reported here (He = 0.153) was slightly lower than what previously reported for other species using SNP panel (0.198-0.220 in East Atlantic Peacock wrasse [53]; 0.233 - 0.290 in European sea bass [59]; 0.270 – 0.310 in Atlantic herring [60]. We cannot exclude that this is due to the low minor allele frequency allowed in the marker dataset (MAF > 0.005), as previously reported in other studies using the same genotyping approach, where even lower values were recorded, likely due to a threshold MAF of 0.002 used [61].

### 4.2 Outlier allele frequency analysis: convergent adaptation in distant sea bream populations?

Analysis based on outlier SNPs revealed a) a strong subdivision of East Mediterranean wild populations into the two “sub-basins” ION and AEG, and b) similarity between Atlantic and Aegean samples, despite the geographic separation. This is reflected by F_ST_ values calculated with OL dataset, that showed the highest values in the comparisons between ATL and ION samples and highly significant values also between ION and AEG. Within-basin, significant differences were found in the Atlantic between the environmentally different groups ATL1/FRA-1 and ATL3/SPA-2, whose pairwise F_ST_ was not significant when using the entire dataset. These results can be ascribed to environmental differences, as highlighted also by seascape analysis. Separating loci under the effect of varying environmental factors, from those whose variation is only due to demographic factors, is not always easy [62]. The frequency patterns across populations of some OL loci showed allele frequency variations more likely related to geographical distance (e.g. 10524_58), but other OLs seem to reflect the effect of barriers between basins (e.g. 7513_19 and 12615_64). This separation is important if the aim is finding genes involved in adaptation, for which a test based on the correlation between allele frequency and environmental factors is expected to work better [49]. In our case, Bayenv analysis added 18 potential outlier loci to those previously detected with Bayescan and Arlequin, many of which showed correspondence with annotated genes and pointed to two biological processes likely related to adaptation to the environment. While we acknowledge that our functional analysis is limited, the results point toward a real role of these SNPs in the adaptation to the environment and deserve further exploration. Therefore, Bayenv approach resulted useful to detect hidden signature of adaptation despite the reported rate of false positives (around 20-50% [63–65]).

## 5. Conclusions

The results presented in this paper are a pivotal step for the development of traceability and conservation tools for the gilthead sea bream. In addition, in a context of massive aquaculture production of this species, an accurate assessment of the wild genetic variability and structure is fundamental to be able to monitor the potential effects of aquaculture in the future. Sea cage fattening is indeed a source of potential escapees of cultured individuals (often selected for productive traits and/or coming from different areas) into the wild, which can affect the genetic characteristics of the wild counterpart [66]. Our results from Greek time replicates, despite being limited in space, suggested that in the last ten to 15 years the genetic of local wild populations has not been affected by aquaculture escapes, despite escape events have been reported, as also found in other studies of the same area [67]. On the contrary, effective population size was higher in the more recent samples.

Assessing population specific neutral and potentially adaptive genetic traits will allow a proper monitoring of the changes that escapees can produce if they successfully reproduce in the wild. The results of our study, based on samples from the entire gilthead sea bream distribution, described the presence of a genetic population structure of the species in the Atlantic and Mediterranean basins and answered biological questions that will support the management of wild stocks of this economically important species and may be used as a baseline for the assessment of the genetic impact of sea cage aquaculture.

Data for this study are available at NCBI: details to be completed after manuscript is accepted for publication

## Acknowledgements

The project is funded by the 7th Framework Programme for research (FP7) under "Knowledge-Based Bio-Economy - KBBE", Theme 2: "Food, Agriculture and Fisheries, and Biotechnologies" Project identifier: FP7-KBBE-2012-6-single-stage Grant agreement no.: 311920. Also, Dr Gkagkavouzis was funded by a PhD scholarship of Alexander S. Onassis Public Benefit Foundation (GR).

## Supplementary material description

**S1.** Details of the laboratory protocol for library preparation.

**S2.** Table of environmental data used in the seascape analysis and in the correlation analysis of Bayescan.

**S3.** Table of markers retained dafter quality filters and used for the analysis. SNP: Stacks ID of the marker; Outlier/Bayenv: indicates whether a SNP was identified as an outlier by both Arlequin and Bayescan and/or by Bayenv; Inside gene: indicates if the tag mapped inside a specific gene; CDS: indicates if the tag mapped in the CDS region of the gene.

**S4.** DAPC scatterplot of the populations used in this work. The barchart indicates the Discriminant Axes eigenvalue. Circles represent 95% inertia ellipses.

**S5.** Pairwise F_ST_ values calculated using all the SNPs. * when p-value < 0.05; ** when p-value <0.01.

**S6.** Plot of the allele frequencies of the outlier markers and markers identified as correlated to environmental parameter. Populations are ordered according to the four genetic clusters identified. For markers identified by Bayenv, values of the correlated environmental parameter are indicated by blue bars.

**S7.** Pairwise F_ST_ values calculated using neutral (above diagonal) and outlier (below diagonal) SNPs. * when p-value < 0.05; ** when p-value < 0.01.

**S8.** Structure plot for the most likely *k* (=2) using only the neutral markers.

**S9.** Structure plot for the most likely *k* (=3) using only the outlier markers.

**S10.** Table of the GO terms indicating biological processes, cellular components and molecular functions related to the genes found close to the outlier markers.

